# A general model of population dynamics accounting for multiple kinds of interaction

**DOI:** 10.1101/2020.05.22.110916

**Authors:** Luciano Stucchi, Juan Manuel Pastor, Javier García-Algarra, Javier Galeano

## Abstract

Population dynamics has been modelled using differential equations almost since Malthus times, more than two centuries ago. Basic ingredients of population dynamics models are typically a growth rate, a saturation term in the form of Verhulst’s *logistic brake*, and a functional response accounting for interspecific interactions. However intraspecific interactions are not usually included in the equations. The simplest models use linear terms to represent a simple picture of the nature, meanwhile to represent more complex landscapes, it is necessary to include more terms with higher order or analytically more complex. The problem to use a simpler or more complex model depends on many factors: mathematical, ecological, or computational. To address it, here we discuss a new model based on a previous logistic-mutualistic model. We have generalised the interspecific terms (for antagonistic and competitive relationships) and we have also included new polynomial terms to explain any intraspecific interaction. We show that by adding simple intraspecific terms, new free-equilibrium solutions appear driving a much richer dynamics. These new solutions could represent more realistic ecological landscapes by including a new high order term.

## Introduction

In the times of the coronavirus, many news on television and magazines try to explain how the size of the infected population evolves, showing exponential plots of the infected populations over time. These communications try to predict the time evolution of the size of this population in the future. Behind these predictions there is always a differential equation model. These polynomial models have linear terms, but to account for more complex interactions they can add higher order terms, as quadratic, cubic, or even, analytically more complex functions, such as decreasing hyperbolic terms. The problem of choosing a complex or a simple model depends on the balance between properly representing nature and being able to understand the model response. In many cases, the simplest model may be enough to understand the benchmarks in the big picture, but sometimes we need more complexity to represent significant aspects of our problem, and therefore, we need more complex and more difficult models. Finding the balance between simple and complex is a tricky problem, but how simple or complex should the model be? Let’s try to answer this question in a population dynamics problem.

In the study of population dynamics, Lotka^1^ and Volterra^2^ were the first ones to model trophic interactions in order to study predator - prey relationships within two (or more) populations. 

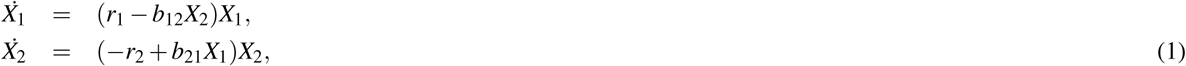

where *b*_*ij*_ terms represent the rate of the interactions between populations *X*_*i*_ and *X*_*j*_ and the *r*_*i*_ represent their effective growth rates. In these equations, signs are incorporated to give a clear meaning to each term, considering all parameters as positive real numbers. This simple model uses a linear term to represent the interaction with the environment and a pairwise second order term to show the antagonistic interaction between the populations of two different species. It was necessary to introduce a higher order term, to represent this interaction.

Although most population dynamics models first dealt with antagonistic relations, mutualistic interactions are widely spread, e.g.^3, 4^. Garcia-Algarra et al.^5^ proposed a logistic mutualistic model. Their formulation was based on writing an *effective* growth rate as the sum of the *intrinsic* growth rate (*r*_*i*_) plus the mutualistic benefit (*b*_*ij*_*X*_*j*_) and, associated with them, to include a saturation term for the whole *effective* growth term. The model was depicted as, 

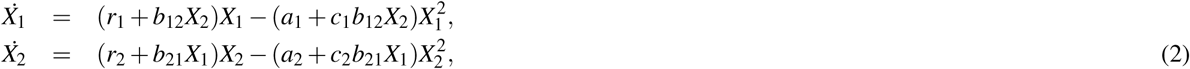

The term 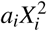 represents the intraspecific competition for resources, and the term 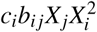 plays the role of saturation for the mutualistic benefit. This model needs to reach a third-order term to prevent the unbounded growth and depicts a well behaved, system, with enough richness to model large ensembles of mutualistic networks and their behaviour.

Other authors have addressed different strategies to introduce the mutualistic interaction. For example, Dean^6^ introduced an exponential dependency on the carrying capacity, *K*, which consequently yields non-linear terms into the equations. To avoid the unbounded growth,^7^ and^8^ proposed restrictions using a type II Holling functional response. This functional leaves the path of introducing a polynomial term with a hyperbolic function.

Nowadays, several studies have focused on adding higher order terms to explain more complex ecological interactions.^9^ studied the influence of interspecific interactions as non-additive density-dependent terms only for competitive communities.^10^ studied the new solutions that a third species adds in pairwise interactions, adding third degree terms with the three different populations, *b*_*ijk*_*X*_*i*_*X*_*j*_*X*_*k*_. It is well-known that an increase in the order of a polynomial term introduces new solutions to the equations, but as^11^ showed these new terms do not always produce viable solutions, furthermore they must be free equilibrium points, and of course the solutions must have an ecological meaning.

Here, we propose a new general model in which any ecological interaction can be included in a simple way. In a first step, we generalise the model proposed by Garcia-Algarra et al.^5^ overcoming the restrictions of the sign of the parameters and, in a second step, we reorganize the intraspecific interactions allowing for both positive and negative interactions, and finally, we introduce a third order term to brake any unbounded pairwise interactions.

## Methods and Materials

We define a new general model. The equation 3 represents the population dynamics of the species *X*_*i*_ driven by an effective growth rate, (first parenthesis in the Eq. 3), and limited by a logistic brake, (second parenthesis in the Eq 3). The view of the model is simple and similar to the original Verhulst idea^12^, where the low order terms represent the increases of the population and the high order terms the brake. The differences with other models are in the terms included in the effective growth rate and logistic brake. The effective growth rate includes the vegetative growth rate, *r*_*i*_, and all density-dependent pairwise interactions, interspecific interactions, *b*_*ij*_*X*_*j*_ (∀*j* = *i*) and intraspecific ones, *b*_*ii*_*X*_*i*_; and the logistic brake includes the logistic term due just to intraspecific competition, *a*_*i*_, the interspecific intraspecific brake, *b*_*ij*_*X*_*j*_*X*_*i*_, and the intraspecific ones, 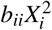.

### A new general model including intraspecific interaction terms

Regarding the mutualistic model (Eq 2), we introduce two differences: First, the parameters of the equation, *r*_*i*_ and *b*_*ij*_ can be positive or negative, representing the different ecological interactions and second, we include the effect of the population in its own effective growth rate just adding the index *j* = *i* in the sum of the interactions terms, so the model can be represented as: 

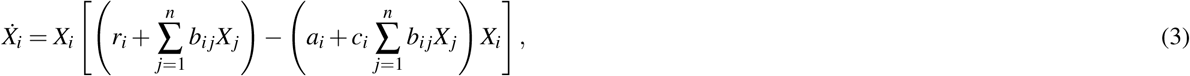

where the subscript *i* runs from 1 to *n*, including the intraspecific interaction (*j* = *i*). With this term we are taking into account the interaction between individuals of the same populations. The new terms yield new solutions and a different phase space. In particular, the inclusion of the term 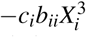 is key for the emergence of new solutions, although there was already a term with the same order in the mutualistic logistic model (Eq 2), 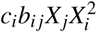. It can be observed in Fig. 13. We explain all details about the number of solutions in appendix

Generally, in the literature of populations dynamics the intraspecific interactions have been introduced only as a logistic brake, *-a*_*i*_*X*_*i*_, representing a growth limit due to resource sharing. In our model, the term, *b*_*ii*_, can represent any kind of intraspecific interaction from beneficial, namely, *cooperation* to harmful interactions, such as *competition* or even *cannibalism*. Even though the logistic term *-a*_*i*_*X*_*i*_ can be seen as the result of intraspecific interactions that limit the growth by resource sharing and it can be included in the interaction term *b*_*ii*_*X*_*i*_, we maintain the separated formulation for the sake of comparison with the equation without this new term.

In fact, there are abundant examples of different intraspecific behaviours in the literature, such as those mentioned above. Cooperation is well known among social and eusocial species^13^ and benefits of cooperative behaviour has been consistently reported, specially for eusocial animals^14^. On the other way, in nature, we can find different types of competition among members of the same population. For example,^15^ reported the aggressive intraspecific behaviour of the Peruvian booby, which attacks their peers not by means of taking their food, but for the sake of being around their nest. In the same way, adult boobies show little tolerance for pigeons that are not from them, pecking them to death. These behaviour is well known for other territorial animals and it conceptually differs from the conventional intraspecific competition for resources.

### Solutions with one population

In general, the equation system (3) cannot be solved analytically. However, the study of only one population can be solved and illustrates the possibilities of the model.

Considering Eq. 3 for only one population. The equation can be written as: 

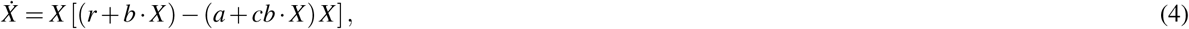

where we have removed the subscripts for simplicity. Stationary points, where 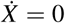 give us the keys to understand the behaviour of the time evolution of the population sizes. The trivial solution, which corresponds to extinction, is *X*^*\**^= 0. Now, the non-trivial stationary solutions can be obtained from the condition: 

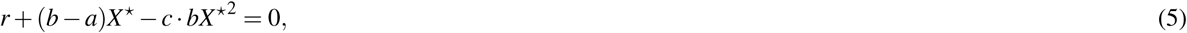

Then, the stationary solutions of Eq. 4 are the extinction and the solutions of Eq. 5: 

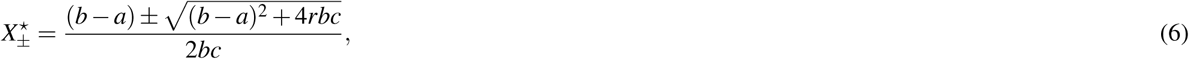

In Ecology, we are only interested in the positive real solutions, generally called feasible solutions. To obtain these feasible solutions in Eq. 6 we need to study several cases, when:

- *r* **and** *b* **have the same sign**. In addition to the trivial solution, *X*\*= 0, in both cases, there is one positive stationary point, which corresponds to the carrying capacity of the population, and the other is negative, which is not a feasible solution.
  – In the case that both parameters are negatives, *r* < 0 and *b* < 0, the positive solution is unstable and the trivial solution is the unique stable solution.
  – In the opposite case, *r* > 0 and *b* > 0,the carrying capacity is the stable solution.
- *r* **and** *b* **have different sign**. The interesting point of having a high order term comes from the possibility of different signs of the parameters. When *r* and *b* have different sign, there are two solutions as long as the condition *c* ≤ (*b a*)^2^*/*4 |*rb*| is fulfilled.
  – If *b* > 0, it is a necessary another condition to obtain a feasible solution that *b* > *a*. Ecologically speaking, this means that the term of intraspecific interaction overcomes the intrinsic growth deficiency and increases the population. in Figure 1b, we plot a case with these conditions. We obtain three fixed points: initial and end points are stable and the intermediate point unstable. This point marks the threshold population; above this value, intraspecific cooperation moves the population to reach the carrying capacity, and below this value, the population goes to extinction.
  – if *b* < 0. In this scenario, the intermediate point is stable and the other solutions are unstable. Consequently, the intraspecific competition produce a a new stationary solution, lower than the carrying capacity. This behaviour has been called as Allee effect^16^. See the example in Figure 1b.

In Fig. 1 we depict on the top, 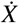 vs. *X* and on the bottom the temporal evolution of the population size, *X* (*t*) vs. *t*. On the left, the growth rate *r* is negative and the intraspecific interaction coefficient *b* is positive. The intermediate stationary solution plays the role of a population threshold because smaller communities will go extinct (Population in orange in Fig. 1 on bottom left), while larger communities will grow to its carrying capacity (population in green in Fig. 1 on the bottom left). On the right, the growth rate *r* is positive and the interaction coefficient *b* is negative. In this case the carrying capacity becomes unstable and the system evolves to the new stable intermediate solution, because of the detrimental intraspecific interaction. (Both populations in Fig. 1 on the bottom right). Two examples of population evolution are plotted in each scenario, where the orange and green dots in the upper plot depict the initial condition of each evolution in the lower plot.

**Figure 1.**
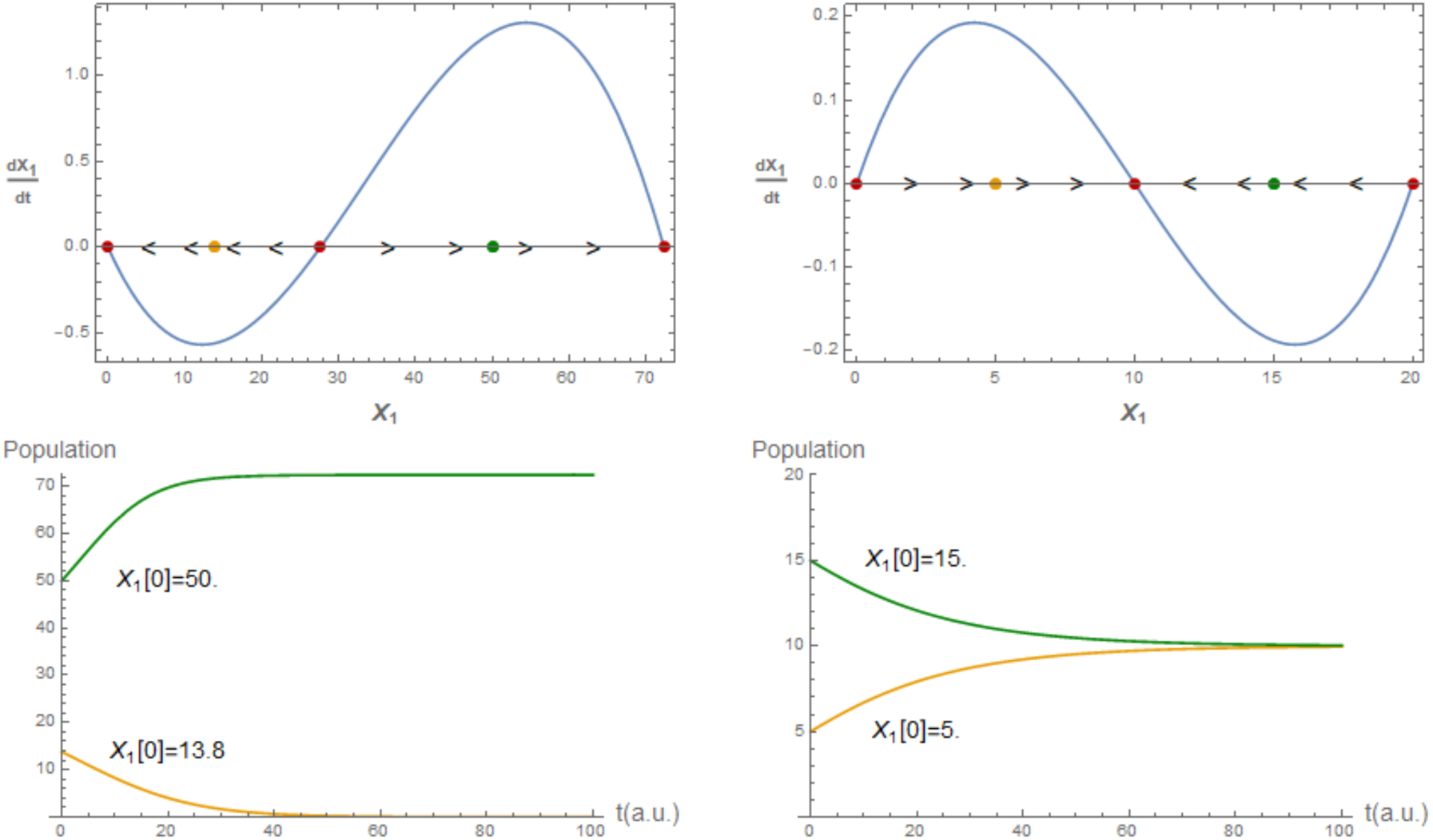
Temporal derivative (up) and population evolution (down) for one population with intraspecific interaction. (Left) Negative growth rate, *r* = −0.1, positive intraspecific interaction, *b* = 0.005 and *c* = 0.005. (Right) Positive growth rate, *r* = 0.1, negative intraspecific interaction, *b* = −0.015 and *c* = 0.05.

### Solutions with two populations

In the case of two populations, the general model is written as: 

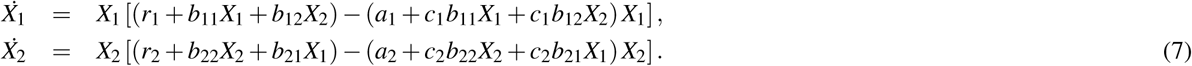

For two populations, we also find the expected trivial solution, i.e. the total extinction 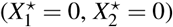, and the partial extinctions, 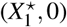 and 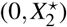, from the equations: 

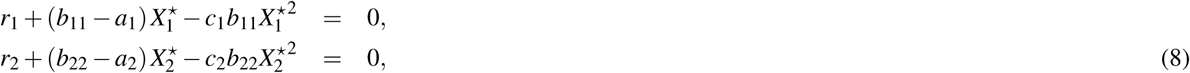

however, as they are second order equations there are two solutions of feasible partial extinctions for each population. The coexistence solutions can be obtained from Eq. 7; these equations can exhibit up to 6 new stationary solutions. Concerning the finite stationary solutions, the intraspecific term makes it more difficult to obtain an analytic expression from the equations: 

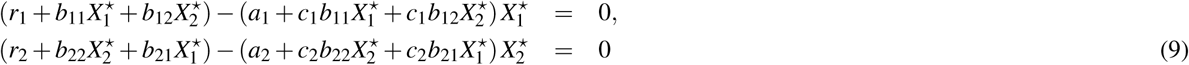

Two out of these six solutions are new free-equilibrium points, due to the new intraspecific terms (details in). Even though we cannot obtain analytic expressions for all solutions, we explored different scenarios by performing numerical simulations with different parameter values. In the next section, we show how the intraspecific interaction changes the phase space of the standard biological interactions.

### Linear stability analysis

In the next section, we explore the linear stability analysis of our system solutions.

#### One population model

To perform the linear stability analysis of the stationary solutions we derive Eq, 4 at the fixed points: 

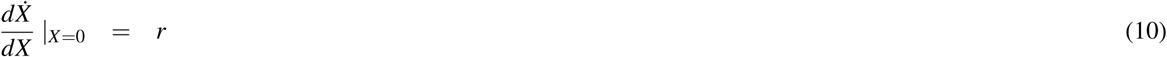

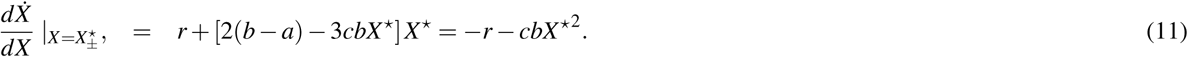

In the trivial solution, the eigenvalue is *λ* = *r* and the unique stable solution is *r* < 0.

According to Eq. 11, the derivative at the (positive) stationary solution *X*^*\**^ will be negative when: 

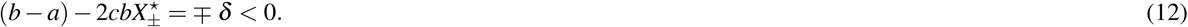

Then 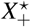 will always be stable and 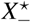 unstable.

When *r* > 0 and *b* > 0, extinction is an unstable solution and population rise to the carrying capacity at 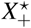, the only positive non-trivial solution. However, for *r* > 0 and *b* < 0, i.e. with intraspecific competition, a new stationary solution emerges, 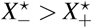. Now the higher solution is unstable and the population only reaches a lower value at the stable point 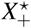. In this case, the negative intraspecific interaction results in a lower carrying capacity.

When *r* < 0, extinction is stable. If *b* < 0 the only positive finite solution is 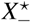, which is unstable. However, when *b* > *a* > 0, a new stable solution, 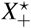, emerges at higher values than 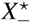. In this scenario, 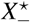 marks the threshold population; above this value, intraspecific cooperation moves the population to reach the carrying capacity, and below this value, the population goes to extinction (see Fig. 1 on the left).

#### Two populations model

The linear stability for the general model (Eq. 3) can be analyzed from the Jacobian matrix at the stationary solutions. Its entries are obtained from: 

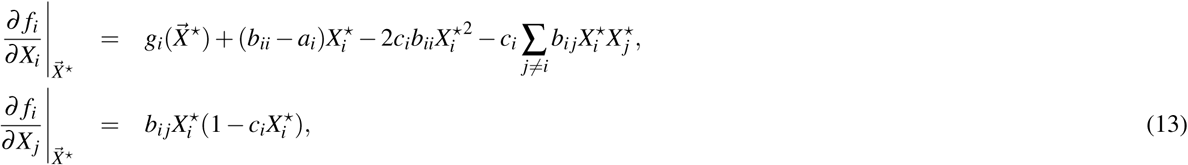

where 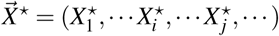 is the vector of the stationary solution.

For two populations, the Jacobian matrix for the total extinction is: 

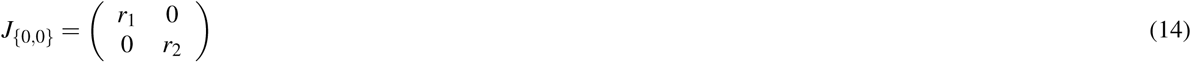

whose eigenvalues *λ*_1_ = *r*_1_ and *λ*_2_ = *r*_2_ are negative when both growth rates are negative.

For the partial extinctions, the Jacobian matrix reads: 

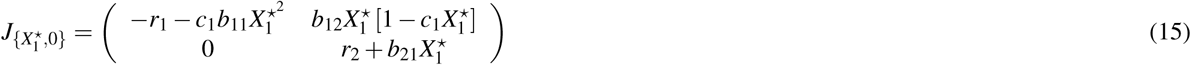

As expected, this Jacobian matrix is almost the same as the matrix for the logistic-mutualistic model (see Appendix A in^5^) but the first entry includes the intraspecific interaction term 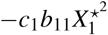. This new term makes the partial extinction to be stable when the intraspecific interaction is positive, *b*_11_ > 0. The same is stated for the symmetric solution 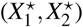.

And for the non-trivial solution 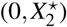 the Jacobian matrix is written as: 

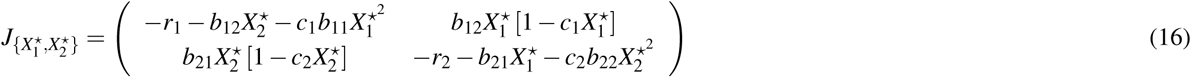

In this case both diagonal entries include the intraspecific term with a negative sign. This means that a positive intraspecific direct interaction enhances the stability of this stationary solution, while a negative intraspecific direct interaction contributes to destabilize it.

A qualitative study of the linear stability can also be made by analyzing the null clines. Solving the null clines, *f*_1_(*X*_1_, *X*_2_)= 0, we obtain two solutions, *X*_1_ = 0 and the expression: 

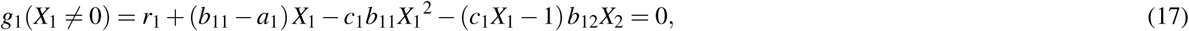

or writing *X*_2_ in terms of *X*_1_, 

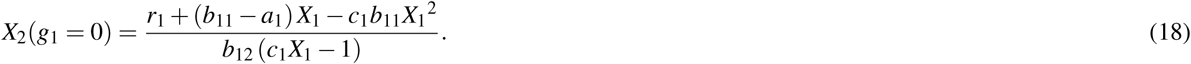

This expression presents a discontinuity at *X*_1_ = 1*/c*_1_, and at *X*_2_ = 1*/c*_2_ for the *f*_2_-null cline. At this discontinuity, the growth rate of species-1 takes the value, *g*_1_(*X*_1_ = 1*/c*_1_) = *r*_1_ − *a*_1_*/c*_1_, independently of *X*_2_ (and the same for *g*_2_(*X*_2_ = 1*/c*_2_)). The condition for a bounded growth leads to *c*_1_ ≤ *a*_1_*/r*_1_, and, as in Verhulst’s equation, this parameter, 1*/c*_1_, plays the role of the carrying capacity. With the same condition for species-2, i.e., *c*_2_ ≤ *a*_2_*/r*_2_, we may define a rectangle limited by *X*_1_ = 0, *X*_1_ = 1*/c*_1_, *X*_2_ = 0, and *X*_2_ = 1*/c*_2_ in whose boundary the flux vectors never point out of the rectangle, and therefore the growth is bounded.

Figure 2 depicts the bounding rectangle limited by the axis and the dashed lines 1*/c*_1_ and 1*/c*_2_. On the left, the conditions *c*_1_ ≤ *a*_1_*/r*_1_ and *c*_2_ ≤ *a*_2_*/r*_2_ are fulfilled and the flux lines are pointing inside the rectangle. On the right, the conditions are no longer satisfied but one stable solution is located outside the rectangle, allowing some flux lines to go out. The asymptotic behaviour of the null cline at *X*_1_ = 1*/c*_1_ has changed and now it rises to infinity.

**Figure 2.**
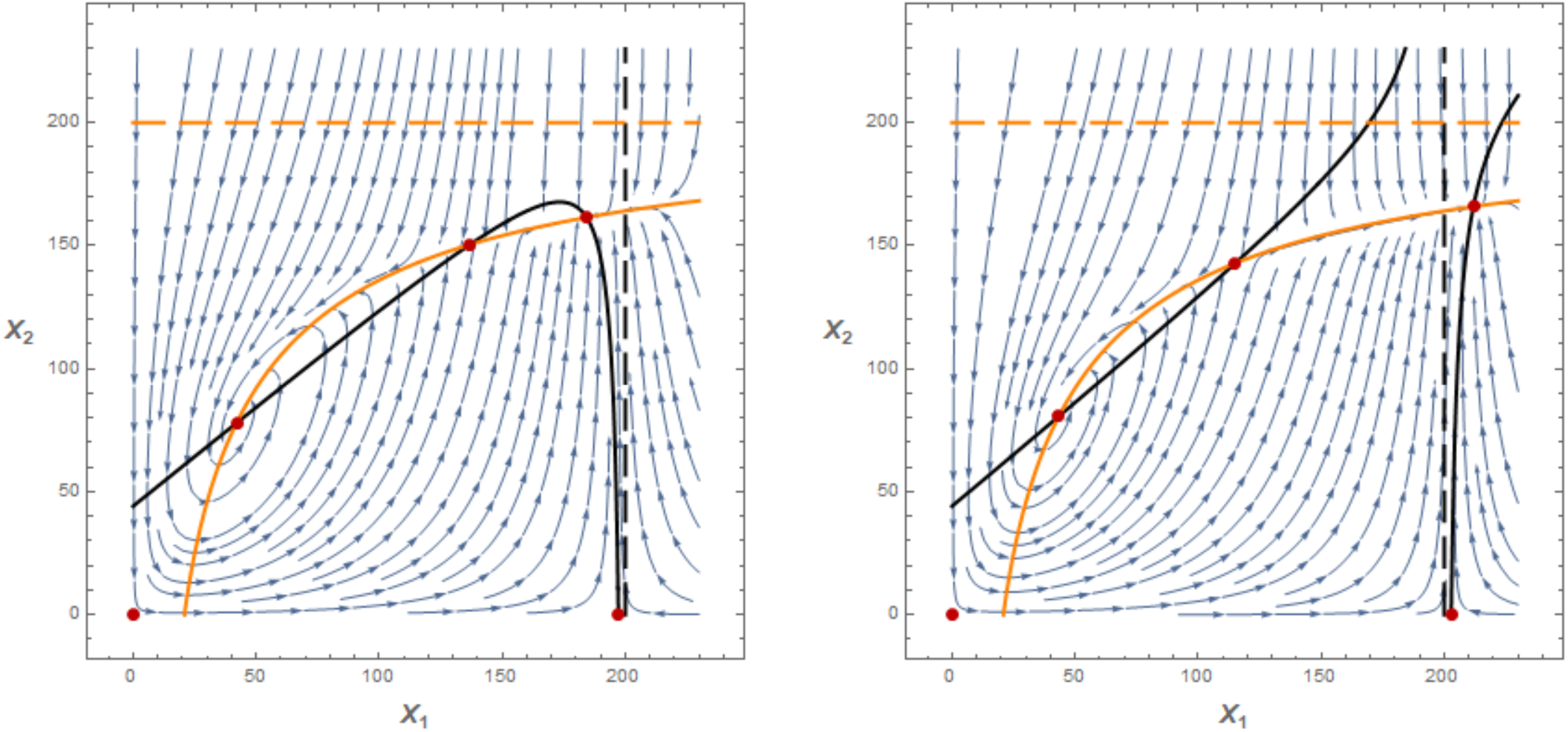
Null-clines and phase space for an antagonistic system where both populations cooperate intraspecifically. Dashed lines represent *X*_*i*_ = 1*/c*_*i*_ while solid lines are the null-clines (orange for *X*_2_ and black for *X*_1_). On the left, *a*_1_ = 0.0008 fulfils the condition *a*_1_ > *c*_1_ *· r*_1_ and flux lines inside the rectangle do not point out of this region; on the right, *a*_1_ = 0.0007 < *c*_1_ *· r*_1_ and some flux lines go out of the rectangle. Parameters: *r*_1_ = 0.15, *r*_2_ = −0.15, *b*_11_ = 0.0028, *b*_12_ = −0.0034, *b*_21_ = 0.0072, *b*_22_ = 0.0005, *a*_2_ = 0.00075, *c*_1_ = *c*_2_ = 0.005.

The intersection of both null clines define the stationary solutions. As the expression Eq. 18 is non linear, there can be several solutions inside the rectangle. This allows more than one stable solution inside this area, separated by saddle points. As an example, Figure 2 shows the intersections of null clines (black lines for *X*_1_ and orange lines for *X*_2_) as red points; two of them are stable stationary solutions, separated by a saddle point. In this example for a predator-prey system, the phase space shows the typical solution of a stable spiral (at *X*_1_ = 42, *X*_2_ = 79) and a new stable node at higher population of predator and prey(at *X*_1_ = 200, *X*_2_ = 164). Note that even though *a*_1_ does not fulfil the condition *a*_1_ ≤ *c*_1_*· r*_1_, in this example, the system is also bounded and stable outside the rectangle. Finally, the same study can be done for *N* species. For every species the value *X*_*i*_ = 1*/c*_*i*_ can define a threshold for the initial population for which the flux trajectories never go outside the N-dimensional rectangle. In this case free-equilibrium solutions will be harder to obtain, however, the Jacobian at these points will have a similar expression (see).

### Solutions with three populations

Ecological complexity increases with species number. Just as a little example we show in this section how the intraspecific interaction can change the outcomes in a 3-species predator-prey system. We show how a positive coefficient in the intraspecific term of the prey-1 avoid the extinction. Figure 3a shows the time evolution of three populations, two preys and one predator; the cooperation coefficient in prey-1 (*b*_11_ = 0.001), even smaller than the interspecific coefficient (*b*_13_ = - 0.004), changes the initial outcome resulting in a stationary population for prey-1 and predator, and the extinction of prey-2.

**Figure 3.**
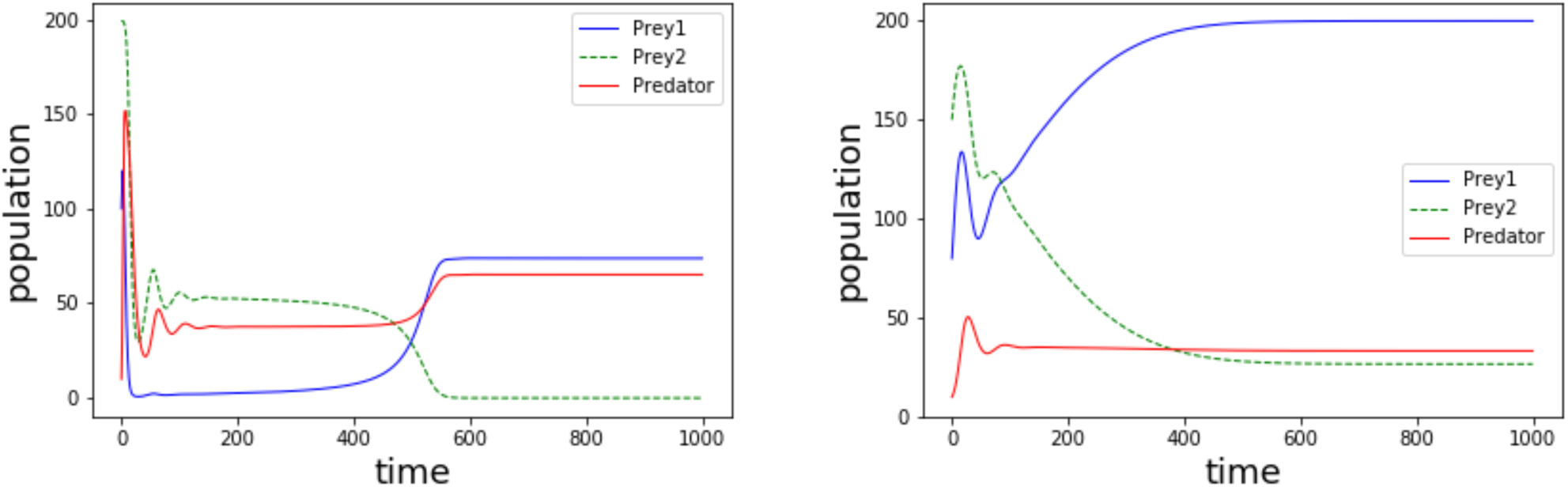
Population evolution in a predator-prey system with two preys. (a) Prey-1 with cooperation *b*_11_ = 0.001 (*r*_1_ = *r*_2_ = 0.15, *r*_3_ = −0.15; *b*_13_ = *b*_23_ = −0.004, *b*_31_ = *b*_32_ = 0.004) ; (b) Predator with competition, *b*_33_ = −0.0005 (*r*_1_ = *r*_2_ = 0.15, *r*_3_ = −0.15; *b*_13_ = −0.004, *b*_23_ = −0.0045, *b*_31_ = *b*_32_ = 0.001)

For the case of negative intraspecific interaction, we show another predator-prey system, with two preys and one predator. In this example the intraspecific coefficient of the predator (*b*_33_=-0.0005) allows both preys to survive at higher populations (Figure 3b); the three populations exhibit initial oscillations until they reach a stationary population, however the difference in the interspecific coefficient (*b*_13_ = − 0.004 and *b*_23_ = − 00045) makes the prey-1 stationary population to be higher than that of prey-2.

## Results

Here we show the great variety of scenarios of ecological interactions that this general model is capable of producing. The aim of this section is to show the great richness of the model, but it is not an exhaustive study of the parameters. We show some examples of the solutions that the intraspecific interaction provide to the populations model with two populations. Since exploring all the possible combinations of signs and ratios among the parameters would be unmanageable and redundant, we only show some interesting cases. For all the figures shown in this section, we have varied the parameters in the effective growth rate, *r*_*i*_, *b*_*ii*_, and *b*_*ij*_ and we have set the limiting parameters *a*_1_ = *a*_2_ = 0.00075 and *c*_1_ = *c*_2_ = 0.005.

### Antagonism

The solutions of the classical predator-prey model are modified when intraspecific interactions comes into play. In our following examples, we have *X*_1_ as the prey and *X*_2_ as the predators. We only show obligate predation, since the facultative case only offers a minor change.

#### The effect of cooperation among prey

The predator-prey system without any intraspecific interaction has only two free-equilibrium solutions: one convergent spiral and one unstable solution, located at the carrying capacity of the prey. The addition of cooperation among the population of preys can generate a new stable solution. Besides the well-known oscillatory solution we may find a new stable node at high population values, separated by a saddle point. Figure 4, left, shows the phase space and trajectories (with the stationary solutions as red points) for a predator-prey system when preys (*X*_1_) exhibit positive intraespecific interaction (*cooperation*). Phase trajectories keep around the stable spiral for low populations, however the saddle point define a new basin towards the new stable solution for high population values (note that the intraespecific parameter, *b*_11_ = 0.0028, is lower than the absolute value of the interspecific parameter, *b*_12_ = − 0.0036).

**Figure 4.**
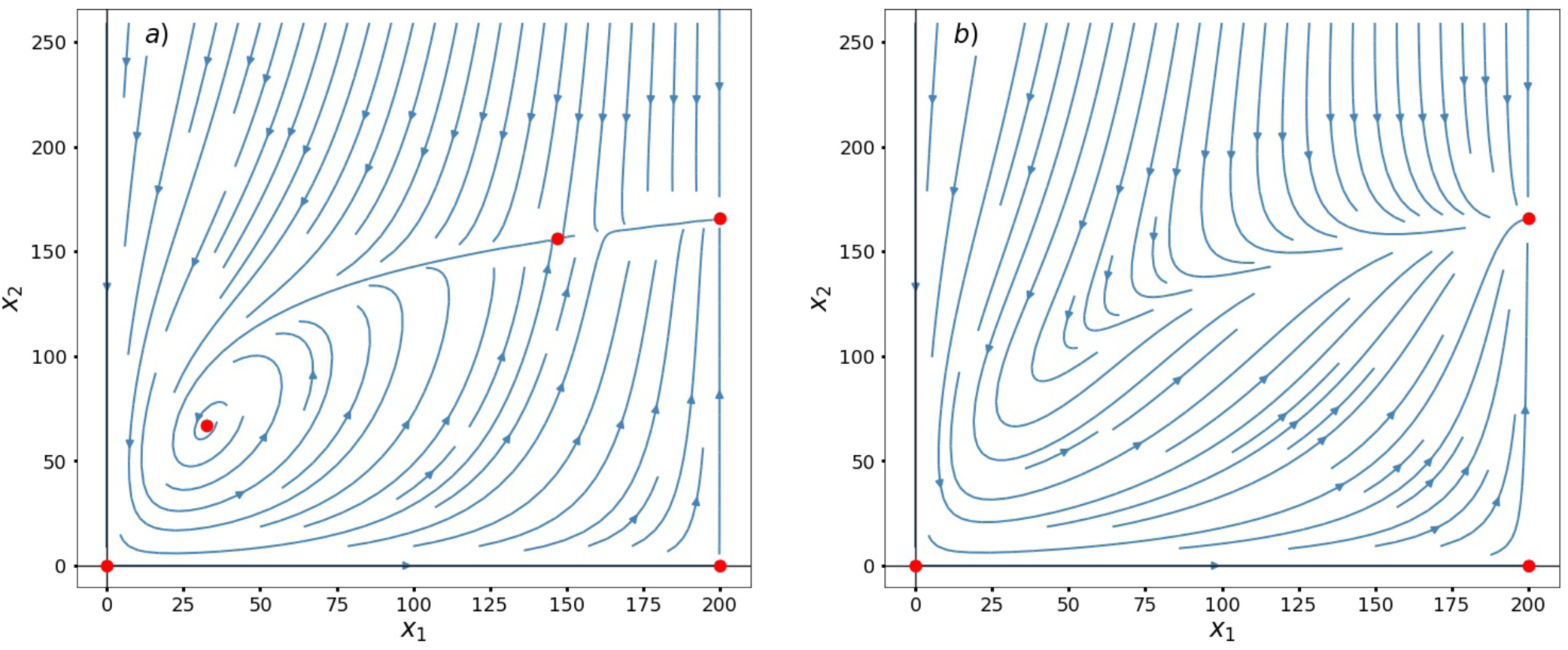
Phase space and trajectories for two populations involved in a predator-prey interaction. To the left we have a case with a lower *b*_11_ = 0.0028 than to the right, *b*_11_ = 0.0035. Cooperation among prey allows a new intermediate solution, which is unstable, and acts in the same way that in Fig.1. Also, as greater cooperation decreases the predatory term, the relation may become commensalistic at some point. Here *r*_1_ = 0.15, *r*_2_ = −0.15, *b*_12_ = −0.0036, *b*_21_ = 0.0072.

If the detrimental interspecific interaction becomes less harmful, the original stable spiral may disappear and the only stable solution is the coexistence at the carrying capacity (Figure 4, right, with *b*_12_ = −0.0036).

When the intraspecific interaction is greater than the interspecific interactions, in our example, |*b*_12_| = *b*_21_ < *b*_11_, a new dynamic appears. The spiral becomes unstable and the trajectories go outwards; as this stationary solution is in the repulsion basin of the saddle point the trajectories cannot go out and they will remain in a closed orbit, i.e., in a limit cycle. In Fig. 5 a (with *b*_11_ = 0.0036 and *b*_12_ = − 0.0072, in addition to representing the trajectories and the stationary solutions) we depict 3 initial points (in green, yellow, and orange) corresponding to the time evolution picture shown below. The intermediate solution that appeared due to the cooperation term acts as a threshold between the spiral and the coexistence located at the carrying capacity of the prey, which remains as a stable solution. Now, if we decrease the intraspecific parameter the saddle point moves toward the carrying capacity and all the stationary solutions become unstable, and all the trajectories fall into the limit cycle (Fig 5b). The corresponding time evolution (Fig 5b below) shows fluctuating population for all initial points.

**Figure 5.**
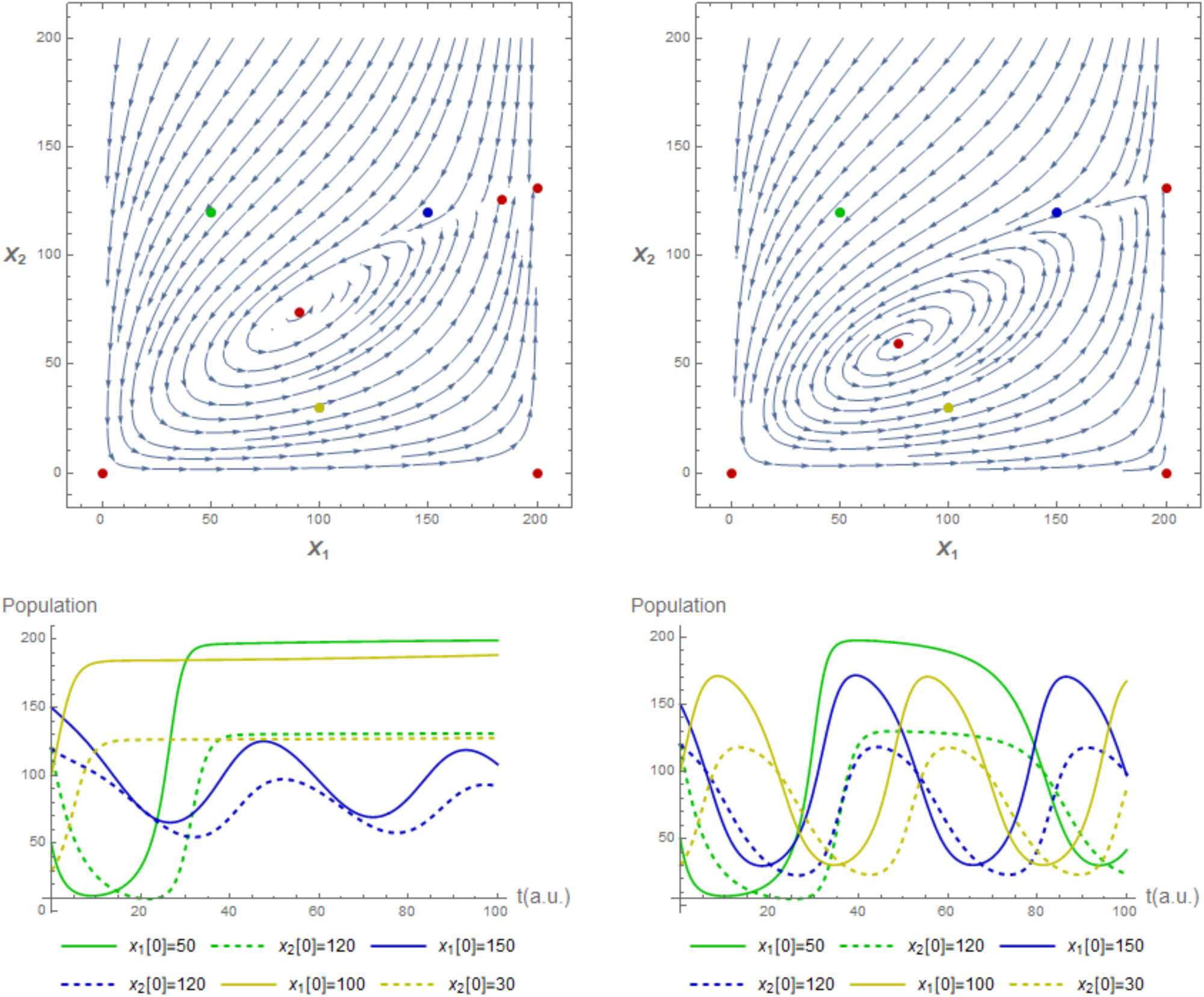
Phase space and trajectories for two populations involved in a predator-prey interaction. We show here a special case where the coexistence spiral solution diverges and become unstable. When that happens, a limit cycle appears. To the left, we have a case with smaller predation, i.e. *b*_12_ = − 0.006435 than to the right, where *b*_12_ = − 0.0072. In both cases, *b*_11_ = *b*_21_ = 0.0036, which means that both populations benefit the same from population *X*_1_ but the predatory effects of *X*_2_ on *X*_1_ are stronger on the right. For greater cooperation values the intermediate solution might even disappear, as it is shown on the right. The green, blue and yellow dots in the phase space mark the initial conditions of the simulations located below. Here *r*_1_ = 0.15, *r*_2_ = −0.15

#### The effect of cooperation among predators

In Fig. 6 we show the effect of the intraspecific interactions only on predators. As in the previous case, without any intraspecific interaction, the system have only two free-equilibrium points, one convergent spiral and one unstable solution, located at the carrying capacity of the prey. The addition of cooperation among predators can generate a pair of new solutions, both of them corresponding to partial extinctions of prey. The effect is the same that we showed for one population in Fig.1 but acting on the predator axis. Thus, cooperation among predators introduces a similar effect of facultative predation. We tested two different values of predators cooperation parameter *b*_22_ to see its direct influence. Although, in both cases the cooperative term is greater than predation, i.e. *b*_21_ < *b*_22_, we can see that at lower values of cooperation almost no effect is notable, but at greater values, two partial extinction of prey appear, one stable and one unstable, a saddle-node bifurcation. This allows predators to survive without preys when cooperation reaches a certain limit. In Fig. 6, to the left, we have the case in which cooperation is weaker and, to the right, the case in which is mildly stronger. The coexistence located at the carrying capacity of the prey remains unstable.

**Figure 6.**
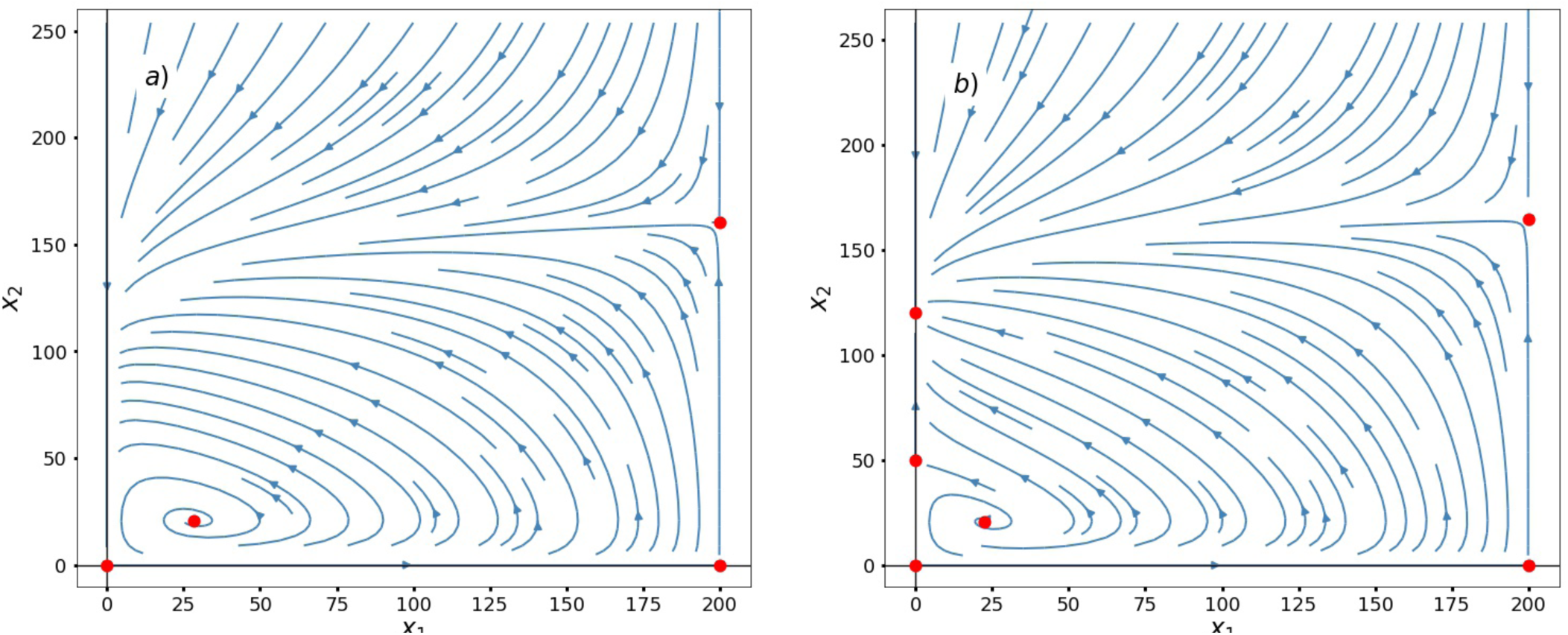
Phase space and trajectories for two populations involved in an antagonist interaction. To the left we have a case with a lower *b*_22_ = 0.004 than to the right, *b*_22_ = 0.005. Cooperation among predators allows two new partial extinctions of prey, one stable and one unstable, in the same way that in Fig.1 but on the predators axis. The coexistence located at the carrying capacity of the prey remains unstable. Here *r*_1_ = 0.15, *r*_2_ = −0.15, *b*_11_ = 0, *b*_12_ = −0.0072, *b*_21_ = 0.0036.

### Competition

In the case of competition, the principle of competitive exclusion stands that the stable solution is the partial extinction but, if interaction parameters are weak, another feasible stable solution is a coexistence point^17^. However, by including intraspecific interactions, the coexistence could become stable for higher or lower values of the interspecific interaction parameters. For a range of positive intraspecific parameters, partial extinctions and the total carrying capacity could be stable at the same time. Adding a positive intraspecific interaction term (*cooperation*) in one species may induce a new saddle point, defining two basins, one towards partial extinction of this species and the other one to the system carrying capacity. When cooperation occurs in both species these two saddle points and the origin define a central attraction basin towards the system carrying capacity, meanwhile outside this basin the system evolves towards one species extinction as stands in the principle of competitive exclusion (see Fig. 7a). When we have negative intraspecific parameters, the carrying capacity becomes unstable, and the only stable solutions are the partial extinctions; however, due to the intraspecific interaction these points occur at a population below its carrying capacity. Both effects can be seen as consequences of intraspecific cooperation and competition in the same way as for one population in Fig. 1. Cooperation induces new solutions as partial carrying capacities and intraspecific competition as partial extinctions.

**Figure 7.**
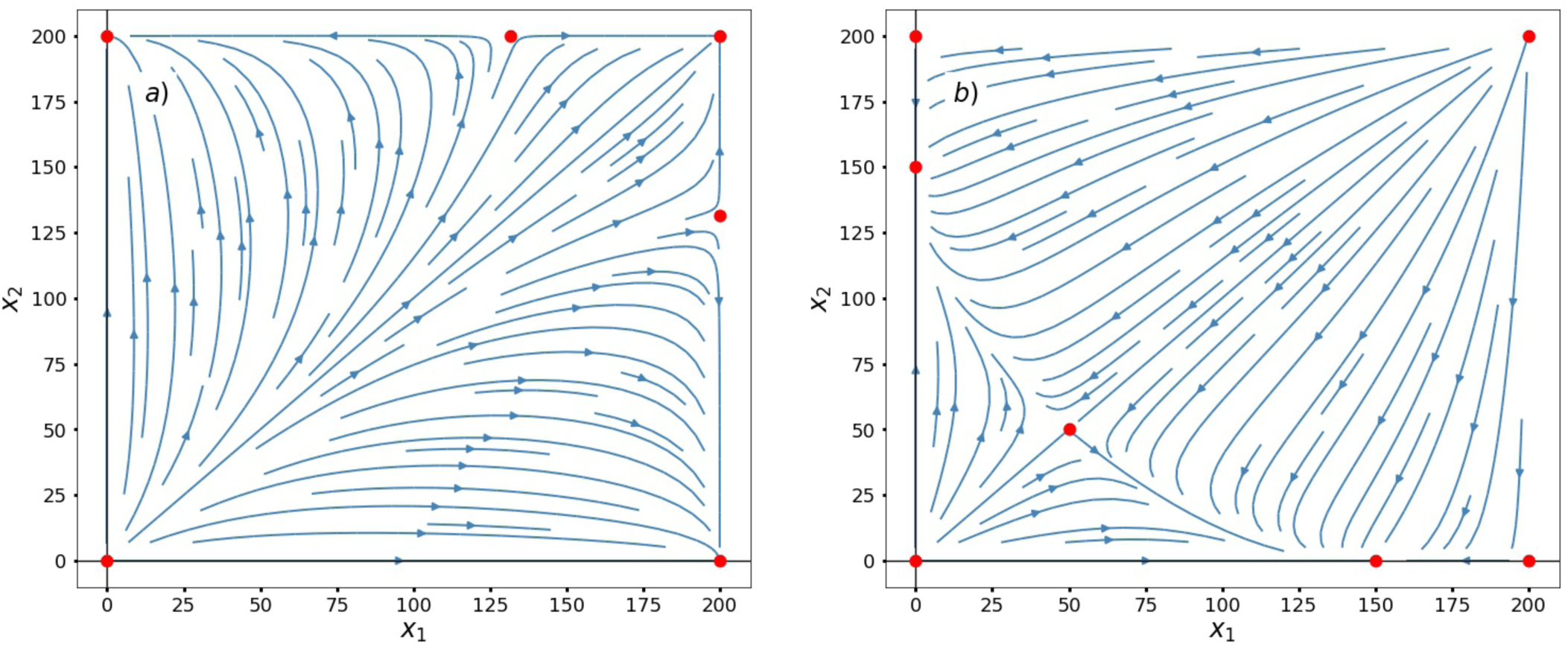
Phase space and trajectories for two populations involved in competition with positive intraspecific interaction. We used two different combinations of *b*_11_ = *b*_22_, to see the influence of intraspecific cooperation and competition. To the left, *b*_11_ = *b*_22_ = 0.0019 and we have the case in which both populations are cooperative and two new solutions appear together with a basin towards the carrying capacity of the system. To the right, *b*_11_ = *b*_22_ = − 0.001 and we have the case in which both are competitive. Noting that when both populations are cooperative, partial carrying capacities appear and they are both unstable. And when both populations are competitive, partial extinctions appear instead, although stables and below the carrying capacities. Here *r*_1_ = *r*_2_ = 0.15, *b*_12_ = *b*_21_ = −0.002.

### Mutualism

The logistic mutualistic model exhibits, in addition to the total and partial extinctions, two feasible finite solutions (^5^): the larger one corresponds to the case where both populations reach their carrying capacities and the lower one is a saddle point that allows us to define a survival watershed. By adding intraspecific interactions, new partial extinctions and carrying capacities could appear.

#### Obligate-obligate mutualism

For the sake of simplicity we only expose the case of equal sign in the parameters for both species, i.e. *r*_1_, *r*_2_ < 0 and *b*_12_, *b*_21_ > 0. In Fig. 8 we show the phase space for two populations involved in a mutual obligatory mutualism with two different values of the cooperation coefficients, *b*_*ii*_. On the left, with weak cooperation the phase space exhibits two free-equilibrium points: the stable carrying capacity and a sadle point defining a *survival watershed*, as in^5^. However, with strong intraspecific interaction (on the right) four new unstable solutions can appear: two saddle points and two unstable fixed nodes, corresponding to partial extinctions. As in the case of one population (see Fig. 1), the new saddle points are the thresholds. Whenever a population is higher than this threshold, it will never go extinct. The total extinction basin is limited by the curve passing through the non-trivial saddle point and these new unstable fixed nodes.

**Figure 8.**
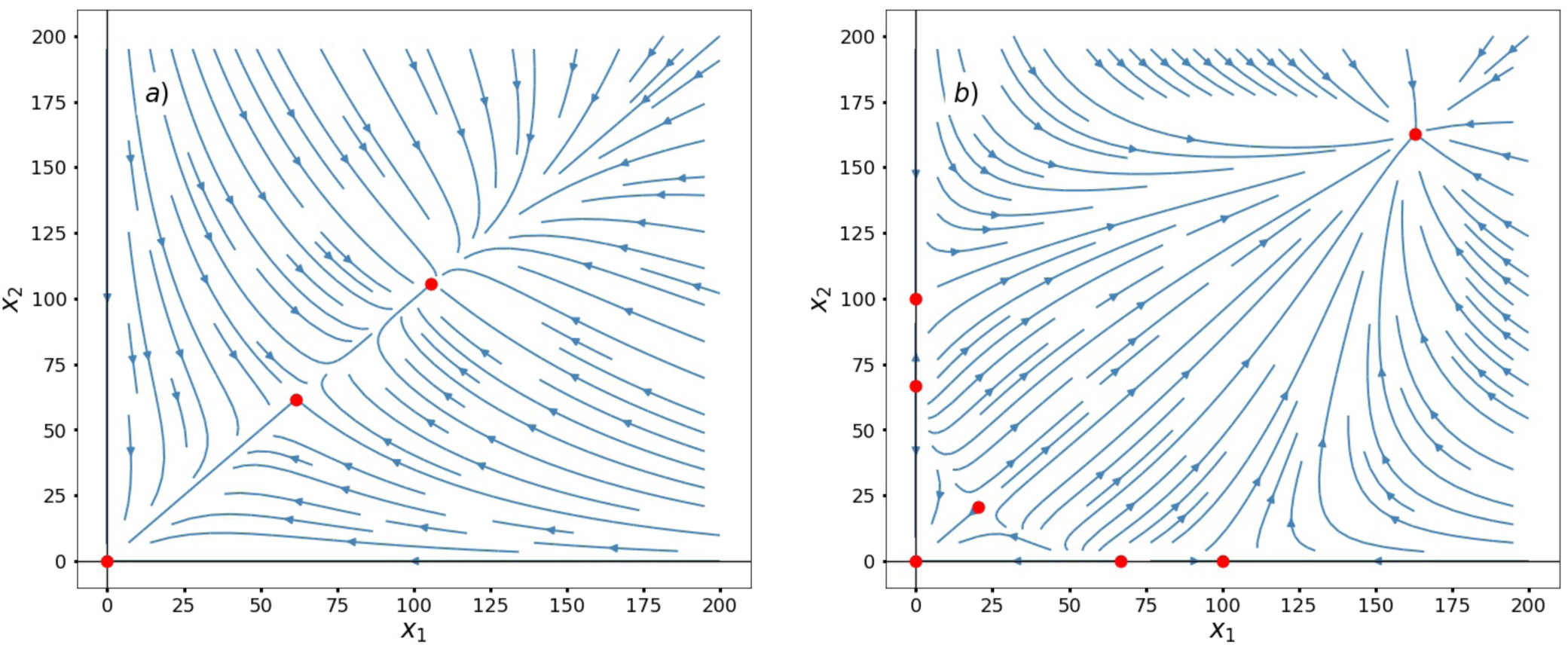
Obligate-obligate mutualism with cooperation in two populations. To the left, we have the case where *b*_11_ = *b*_22_ = 0.0001, which means that intraspecific cooperation is lower than mutualism. To the right, we have *b*_11_ = *b*_22_ = 0.0045, which means that both intraspecific cooperation and mutualism weight the same. Here *r*_1_ = *r*_2_ = −0.15, *b*_12_ = *b*_21_ = 0.0045.

On the other hand, when mutualistic species exhibit negative intraspecific interactions, as in Fig. 9, the stable carrying capacity moves toward the saddle point (on the left). And eventually, when this negative term is high enough, these two solutions collide and total extinction remains as the exclusive stable stationary solution (on the right).

**Figure 9.**
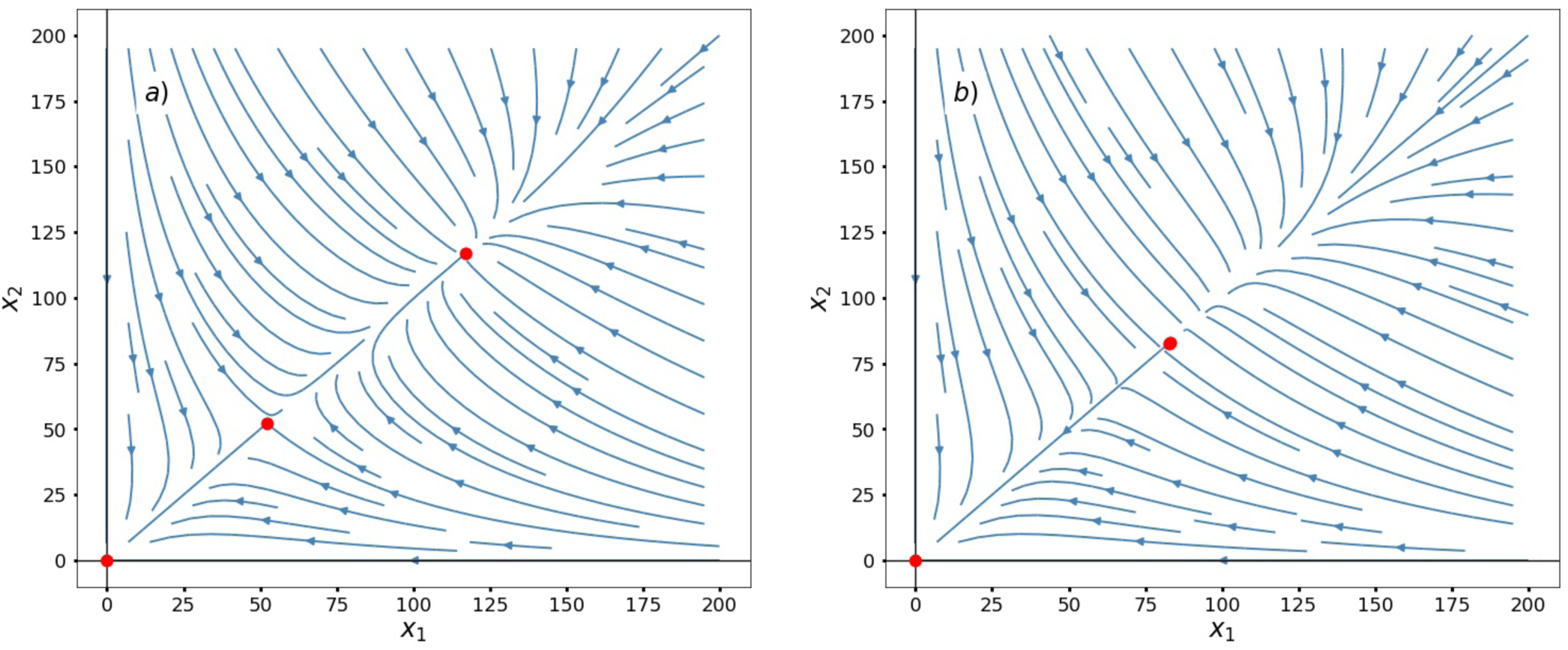
Obligate-obligate mutualism with competition in two populations. To the left, we have the case where *b*_11_ = *b*_22_ = −0.0001, which means that intraspecific competition is lower than mutualism. To the right, we have *b*_11_ = *b*_22_ = −0.00062868, which means that intraspecific competition has stronger effects than mutualism. Here *r*_1_ = *r*_2_ = −0.15, *b*_12_ = *b*_21_ = 0.005.

In the case of one cooperative population and one competitive population the system exhibits this asymmetry and, again, a new saddle point in the cooperative population axis set a survival threshold. Above it, the system always evolves towards the coexistence solution and will never go extinct, as it is shown in Fig. 10.

**Figure 10.**
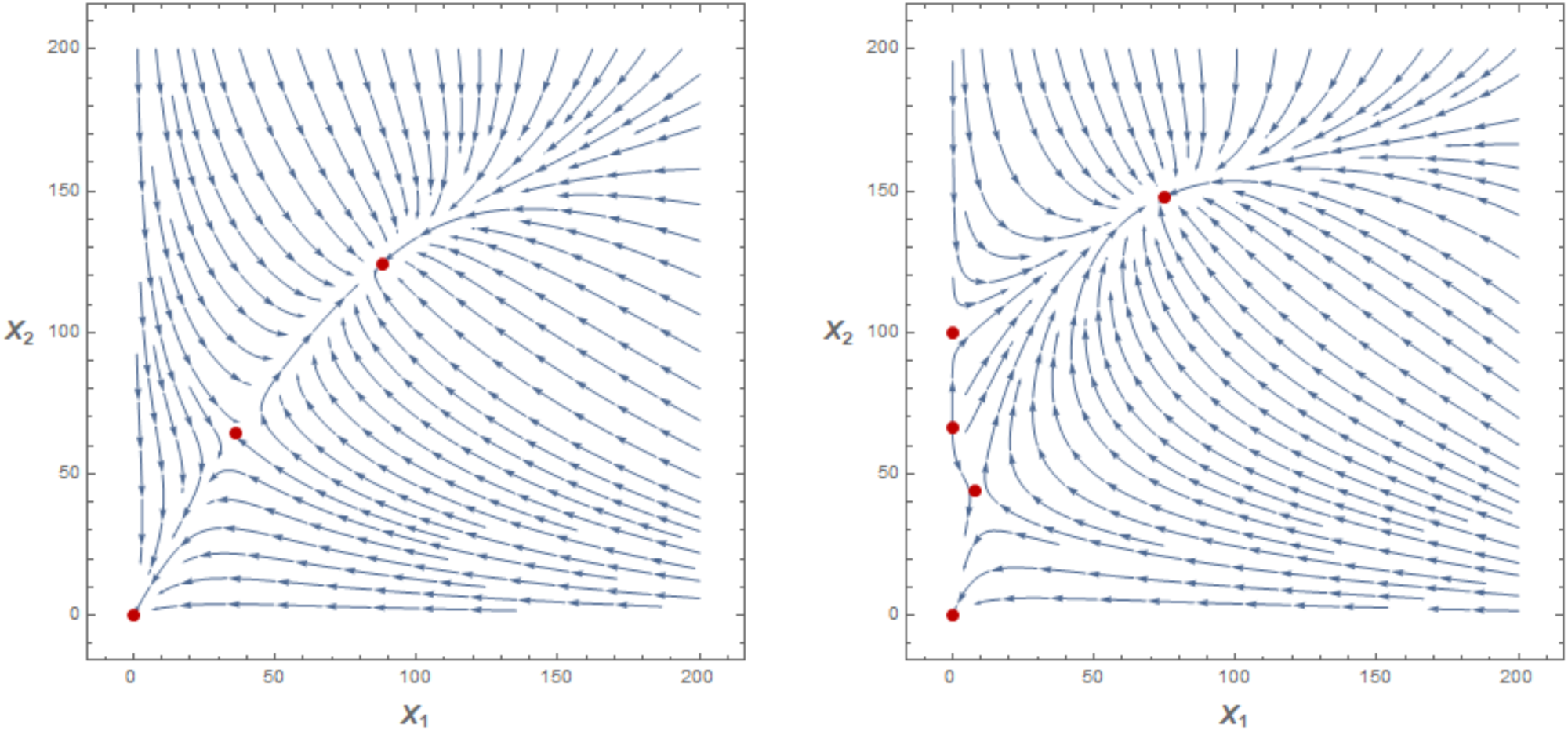
Obligate-obligate mutualism with positive and negative intraspecific interaction.To the left, we have the case where *b*_11_ = −0.002 and *b*_22_ = 0.002, which means that intraspecific competition of *X*_1_ is the same that intraspecific cooperation of *X*_2_ and both interactions weight lower than mutualism. To the right, we have *b*_11_ = −0.0045 and *b*_22_ = 0.0045, which means that intraspecific competition of *X*_1_ weights the same than mutualism and intraspecific cooperation of *X*_2_. Here *r*_1_ = *r*_2_ = −0.15, *b*_12_ = *b*_21_ = 0.0045.

#### Facultative-facultative mutualism

When both growth rates, *r*_1_ and *r*_2_, are positive, total extinction is an unstable solution and the carrying capacity is stable (left plot in Fig. 11). However, when both populations exhibit negative intraspecific interactions the maximum system carrying capacity may become unstable and a new stable finite solution emerges at lower populations (Fig. 11 on the right), as one expects following the one population solution with intraspecific competition (see Fig. 1). In Fig. 11, on the left, the intraspecific interaction generates four partial extinctions as unstable stationary solutions (two saddle points and two unstable nodes). On the right, with higher negative intraspecific interaction, two extra solutions appear as partial carrying capacities, and the total carrying becomes unstable. In this case the system exhibits 9 positive stationary solutions: four saddle points, four unstable points and only one stable solution.

**Figure 11.**
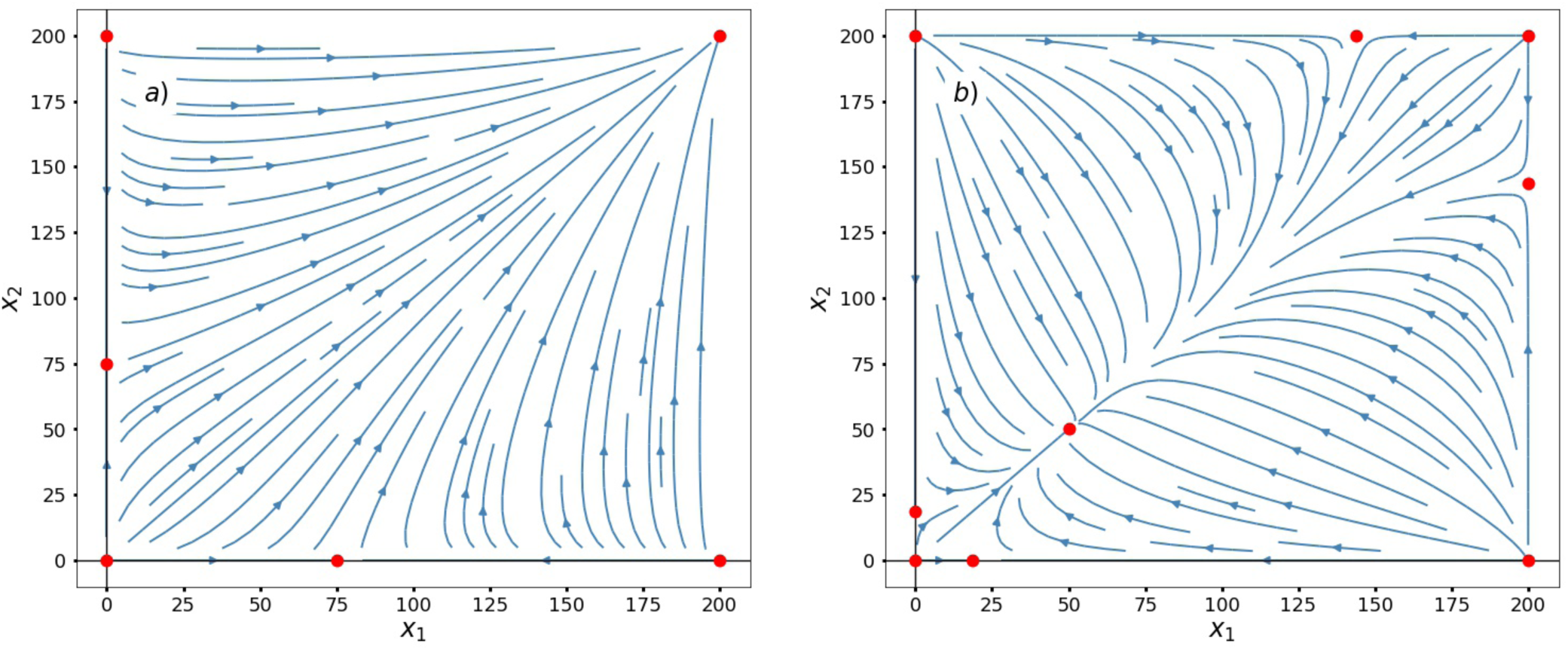
FFM with negative intraspecific interaction.To the left, we have the case where *b*_11_ = *b*_22_ = −0.002, which means that intraspecific competition is weaker than mutualism. To the right, we have *b*_11_ = *b*_22_ = −0.008, which means that intraspecific competition is stronger than mutualism. Here *r*_1_ = *r*_2_ = 0.15, *b*_12_ = *b*_21_ = 0.005.

In the case of facultative mutualism with different intraspecific interactions, one of them beneficial and the other one harmful, the carrying capacity could be reduced for populations with negative intraspecific interaction, while its partner, with positive intraspecific interaction, will grow until reaching its own saturation. Figure 12 depicts this scenario. On the left, competition is weaker than cooperation and the total carrying capacity is the stable stationary solution. On the right, competition is stronger than cooperation and the total carrying capacity becomes unstable. As before, competition only generates unstable a partial extinction, while cooperation pushes the coexistence solution into a transcritical bifurcation.

**Figure 12.**
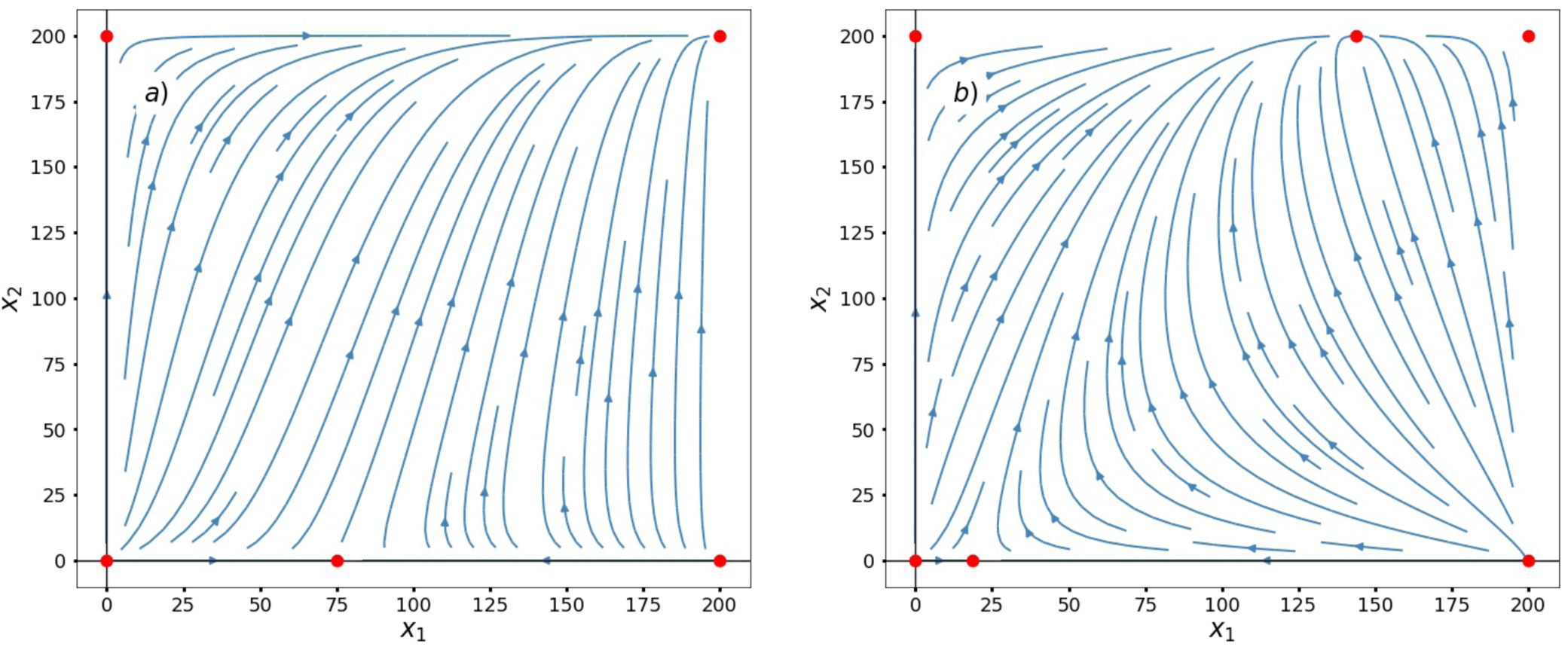
FFM with intraspecific competition and cooperation. To the left, we have the case where *b*_11_ = −0.002 and *b*_22_ = 0.008 and to the right, we have *b*_11_ = −0.008 and *b*_22_ = 0.002. Here *r*_1_ = *r*_2_ = 0.15, *b*_12_ = *b*_21_ = 0.005.

## Conclusions

In the title of the paper we ask how simple should be a population dynamics model. To address the discussion we have introduced the intraspecific interactions in the^5^ model using their same philosophy to include new terms. These appear in the first term of the interaction, representing the effective growth rate, and in the logistic brake to balance the first term. With respect to the previous model, this modification introduce two new terms: 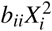 and 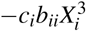, regarding the intraspecific interactions. Furthermore, we have generalized the model allowing the parameters that define the interactions, *b*_*ij*_, to be positive or negative.

In our opinion, the ecological reason to introduce different intraspecific interactions is supported by observations, cooperative and competitive intraspecific interactions are widely known in a wide variety of ecological systems, from social insects to microbial communities. They have been overseen by population dynamics modelling, which mainly focused on interactions with the environment or interspecific interactions (see, for example, the historical sequence developed by^18^).

Furthermore, the cubic term offers an interesting behavior from the mathematical point of view. As^11^ explain that, new high order terms can introduce new free-equilibrium solutions, but it is necessary that these solutions will be feasible, and of course, with a clear ecological meaning. In this way, several authors have used high order interactions to improve the stability or diversity of ecological models. For example,^9^ show the advantages of the high order terms, introducing non-additive density-dependent effects, the authors study the influence of the high order interactions in the competitive communities. Or^19^ show how the high-order interactions increase the stability of the systems. In our model, the term 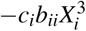 introduces 2 new free-equilibrium solutions (see) that in our opinion can explain ecological situations that were not well explained before with the population dynamics equations.

Delving into the idea of high order interactions,^10^ introduce 3-way or 4-way terms, overcoming the pairwise interactions. These terms are intended to simulate the effect that interactions between species are modulated by one or more species. This idea is inspiring but we believe that simpler models like ours that use polynomial terms and pairwise interaction can still explain many ecological landscapes. Every time that we increase the order of a new term, it is more difficult to define it and their corresponding parameters in the field.

We would like to highlight that the inclusion of the intraspecific terms shows new solutions that could represent more complex ecological landscapes. For example, the case of predator-prey system with positive intraspecific term in the preys exhibit a new solution with a steady state at large populations. This solution could represent the way herds act as a defensive mechanism for preys^20, 21^. Also, large herds of zebras or wildebeest seem to be stable in time; in^22^ the authors presented data of the Kruger National Park, in South Africa, that showed a stable and increasing populations of zebras and wildebeest (more than 10,000 individuals) over a period of twenty years, with an also a more or less stable population of lions (around 400 individuals). Or the effects of intraspecific competition can act as a regulatory mechanism.^23^ showed that intraspecific predation acts in a reinforced way: higher populations decrease the resources available for individuals, reducing their growing rates and promoting smaller and weaker individuals, those are more easily killed or eaten, which increases the per capita food level, both by reducing the population and by satiating the cannibalists.

The main advantage of this general model (Eq. (3)) is that it can be used to describe any ecological regime and that it carries its own saturation mechanism, that avoids the *‘orgy of mutual benefaction’* of^24^.^25^ showed, using a simplified generalized model, studying a nursery pollination system, modelling all the interspecific interactions with the same functional. This allowed a clear interpretation of the parameters of the whole system and an unambiguous way to compare them. Furthermore,^26^ showed that intraspecific interactions in a predator-prey system might lead to diffusion-driven instabilities.

Finally, we would like to venture to discuss some more speculative ideas. Nowadays there are some attempts to model transitions from antagonistic to mutualistic interspecific relationships, limited by the fact that they deal with different mathematical functionals for mutualism and antagonism^27–29^. These models include changes that arise continually from one regime to another, but treating the transition only in a descriptive way. In addition, adaptive changes are modelled through parameter changing systems, where parameters have their own dynamic equations, but these models are still limited to specific ecological regimes, either antagonistic or mutualistic^30–32^. However, if one may adequately define the dynamics of the parameters in a general model of ecological interactions, it may reflect a deeper view of nature, where ecology meets evolution. Thus, by including evolutionary changes in our model, one may be capable of modelling transitions due to mutations and natural selection, which is surely the way how transitions on ecological regimes occur in nature.

## Supporting information

Two sections: Newton's Polytopes and N-dimension Jacobian

## Acknowledgements (optional)

This work was supported by Ministry of Education, Culture, and Sport of Spain (PGC2018-093854-B-I00).

## Author contributions

The authors contributed equally to all aspects of the article.

## Competing interests

The authors declare no competing interests.

## Notes

### Competing Interest Statement

The authors have declared no competing interest.

