## Supplementary material for "A general model of population dynamics accounting for multiple kinds of interaction": Two sections: Newton's Polytopes and N-dimension Jacobian

### Supplementary information

### Quantifying the effect of the intraespecific term

To understand the effect of the intraspecific term in the number of free-equilibrium solutions, we use the technique proposed by[11]. In this appendix, we calculate the number of free equilibrium points in the mutualistic model[5] and in our general model with the intraespecific interaction.

The mutualistic model is depicted, for two species, as,

$$\begin{aligned} \dot{X_1} &= [(r_1 + b_{12}X_2) - (a_1 + c_1 b_{12} X_2)X_1]\, X_1, \\ \dot{X_2} &= [(r_2 + b_{21}X_1) - (a_2 + c_2 b_{21} X_1)X_2]\, X_2, \end{aligned} \tag{19}$$

Following the steps proposed in[11], first we obtain the set of exponents of the monomials of the free-isoclines functions $S_1$ $\{(0,0),(0,1),(1,0),(1,1)\}$ and $S_2$ $\{(0,0),(1,0),(0,1),(1,1)\}$. These set of points allow to plot the Newton polytopes, $P_1$ and $P_2$. Then we need to calculate the Minkowski sum $P_1 \oplus P_2$. And finally we obtain the mixed volume of the Newton polytopes, $M(P_1,P_2)$

$$M(P_1,P_2) = vol_2(P_1 \oplus P_2) - vol_2(P_1) - vol_2(P_2) = 4 - 1 - 1 = 2 \tag{20}$$

consequently, the number of free solution is 2.

Now, we perform the same procedure in our model with the intraespecific term, for two species.

$$\begin{aligned} \dot{X_1} &= \left[(r_1 + b_{11}X_1 + b_{12}X_2) - (a_1 + c_1 b_{11} X_1^2 + c_1 b_{12} X_2 X_1)\right] X_1, \\ \dot{X_2} &= \left[(r_2 + b_{21}X_1 + b_{22}X_2) - (a_2 + c_2 b_{21} X_1 X_2 + c_2 b_{22} X_2^2)\right] X_2, \end{aligned} \tag{21}$$

Now we obtain the new set of exponents of the monomials of the free-isoclines functions $S_3$ $\{(0,0),(0,1),(1,0),(2,0),(1,1)\}$ and
$S_4$ $\{(0,0),(1,0),(0,1),(1,1),(0,2)\}$. These points allow to plot the Newton polytopes, $P_3$ and $P_4$ (see figure 13 ). Then we calculate the Minkowski sum $P_3 \oplus P_4$ and consequently the mixed volume, $M(P_3,P_4)$

$$M(P_3,P_4) = vol_2(P_3 \oplus P_4) - vol_2(P_3) - vol_2(P_4) = 7 - 1.5 - 1.5 = 4 \tag{22}$$

We can stand out that the equation with the intraspecific term, with the same order, introduce 2 new free solutions, with a full ecological meaning, as we can see in Section **??**.

### The Jacobian matrix for N species

For N species, making use of [13], the diagonal terms of the Jacobian matrix at the free-equilibrium points can be written as:

$$J_{ii} = -r_i - \sum_{j \neq i} b_{ij} X_j^\star - c_i b_{ii} {X_i^\star}^2 \tag{23}$$

and the non-diagonal terms:

$$J_{ij} = b_{ij} X_i^\star \left[1 - c_i X_i^\star\right] \tag{24}$$

So, the Jacobian matrix can be written as:

$$J_{\{X_1^\star, X_2^\star\}} = \begin{pmatrix} \ddots & \cdots & \cdots & \cdots & \cdots \\ \cdots & J_{ii} & \cdots & J_{ij} & \cdots \\ \vdots & \vdots & \ddots & \vdots & \vdots \\ \cdots & J_{ji} & \cdots & J_{jj} & \cdots \\ \cdots & \cdots & \cdots & \cdots & \ddots \end{pmatrix}$$

Note that the intraespecific coefficient appears in the Jacobian matrix as a new negative quadratic term.

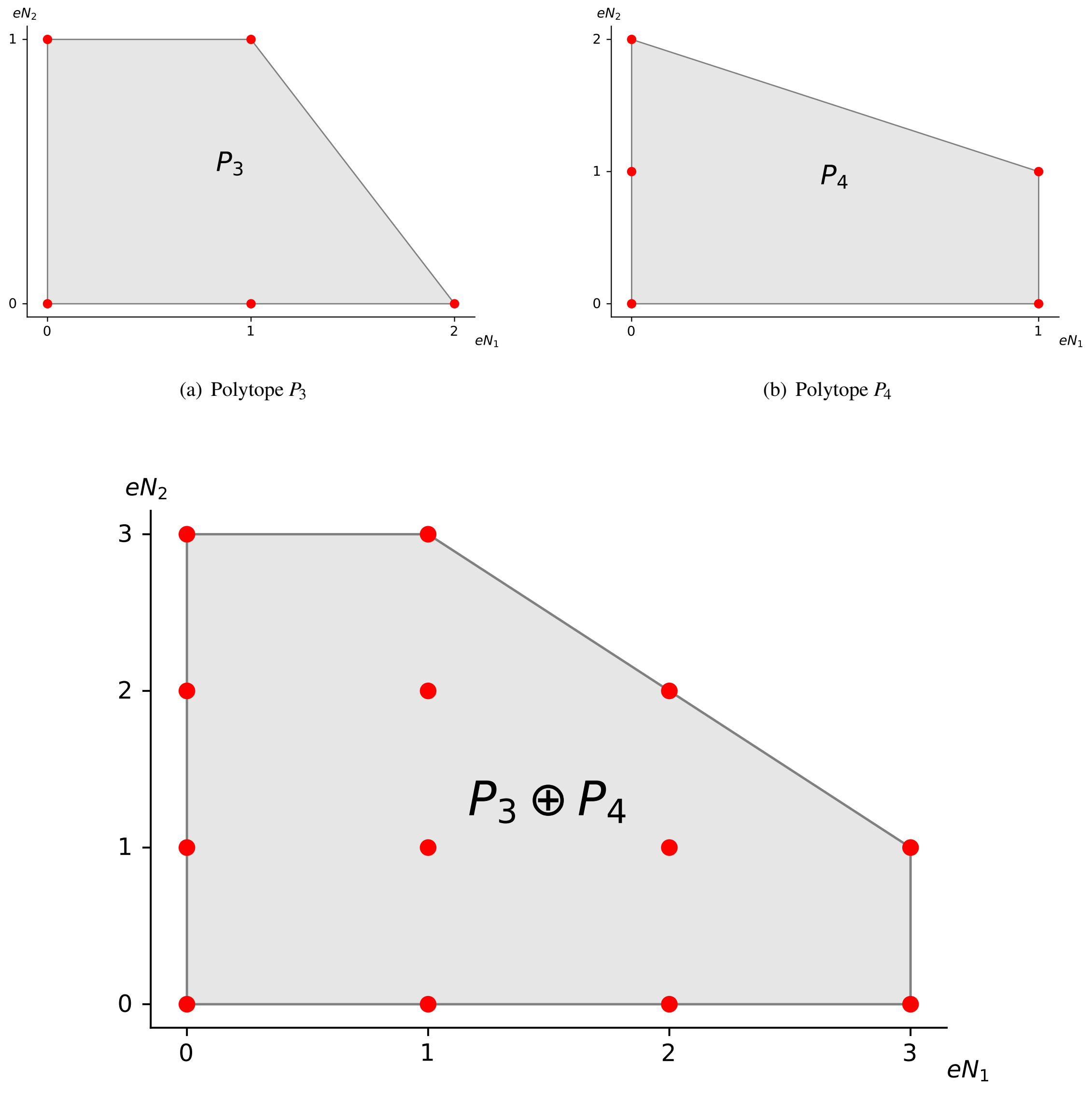


(a) Polytope $P_3$



(b) Polytope $P_4$



(c) Polytope $P_3 \oplus P_4$

**Figure 13.** Mixed volumes of Newton polytopes of the new model with the intraspecific terms. Plots (a) and (b) represent the Newton polytopes of sets $S_3$ and $S_4$ (red points), respectively. Plot (c) represents the mixed volume of the Newton polytope of the Minkowski sum $P_3 \oplus P_4$.
